# *NOR1* and Mitophagy: An Insight into Sertoli Cell Function Regulating Spermatogenesis

**DOI:** 10.1101/2025.02.24.639874

**Authors:** Bhola Shankar Pradhan, Deepyaman Das, Hironmoy Sarkar, Indrashis Bhattacharya, Subeer S Majumdar

## Abstract

The development and differentiation of germ cell (Gc) is supported by the Sertoli cel (Sc). We previously reported that a developmental switch in Sc around 12 days of age onwards in rats. During the process of differentiation of Sc, the differential expression of mitophagy-related genes and its role in male fertility is poorly understood. To address this gap, we evaluated the microarray dataset GSE48795 to identify 12 mitophagy-related hub genes - *Egfr, Bcl2, Ccl2, Mmp2, Igf1, Fgf7, Apoe, Fos, Cxcl12, Ocln, Dcn,* and *Snca*. We identify *NOR1* as a potential mitophagy-related gene of interest due to its strong regulatory association with two hub genes Bcl2 and Fos. To determine the importance of our analysis of finding key mitophagy-related genes, we generated a transgenic rat model with Sc-specific knockdown of *NOR1* during puberty. Adult male transgenic rats were found to be infertile due to impaired spermatogenesis, indicating a critical role for *NOR1* in Sc function. These findings suggest that mitophagy-related genes may play an important role in the differentiation of Sc and thereby, regulate male fertility.

## Introduction

The increase in the cases of male infertility worldwide is a global concern (Agarwal et al., 2021). A recent report suggests the prevalence of infertility is around 17.5% (Vander Borght et al., 2018; Sharpe 2012). The causes of male infertility are diverse and most of the cases are idiopathic in nature (Tournaye et al., 2017; Olesen et al., 2017). Most of them do not respond to any of the conventional modes of therapy, including hormonal supplementations (Bouvattier et al., 2011; Irvine et al., 1998).

The development and function of Sertoli cells (Sc) is important for the process of spermatogenesis, which regulates the division, differentiation and survival of the male germ cells (Gc) into sperm (Griswold, 1998). The proper function of Sc is critical for male fertility. Upon maturation, Sc supports the robust process of spermatogenesis and in many patients with idiopathic male infertility, the maturation of Sc is impaired (Hai et al., 2014). Thus, the understanding of the molecular mechanisms regulating the process of Sc maturation is critical for defining the causes of male infertility.

Mitochondrial dysfunction is contributing significantly to the rising cases of male infertility (Tesarik et al., 2023; Sarkar et al., 2025). The proper function of mitochondria is regulated by several quality control mechanisms inside the eukaryotic cells, such as mitochondrial fusion and fission; and mitophagy (Youle and Narendra, 2011). Mitophagy is the removal of the excessive mitochondria or damaged mitochondria that are beyond repair (Pickles et al., 2018; Mizushima, 2007; Mizushima et al., 1998; Tsukada and Ohsumi, 1993). During the process of mitophagy, the microtubule-associated protein 1A/1B light chain 3 (LC3) is recruited to the autophagosomal membrane, further binding to the damaged mitochondria expressing mitophagy receptors. Later on, mitophagosomes fuse with lysosomes for their degradation (Kuma et al., 2017; McWilliams et al., 2016; Sun et al., 2015). This process of outer membrane fusion of mitochondria is mediated by mitofusins MFN1 and MFN2 and the fusion of inner membrane is mediated by the dynamin-like 120 kDa protein (OPA1). The dysfunction of MFN1, MFN2 or OPA1 leads to various diseases due to compromised OXPHOS activity (Chan, 2020). Moreover, the cascade of events involving phosphatase and tensin homolog (PTEN) induced kinase I (PINK1), a serine/threonine kinase and E3ubiquitin ligase Parkin activates the ubiquitin proteasome system (UPS) (Chan et al., 2011; Rakovic et al., 2019) and recruits autophagosomes for mitophagy (Sekine and Youle, 2018).

In many male infertile patients, various mutations in mtDNA have been reported (Baklouti-Gargouri et al., 2014; Carra et al., 2004; Demain et al., 2017). The mutations in the mitochondrial polymerase gamma (POLG) contribute to male infertility (Demain et al., 2017; Luoma et al., 2004). The mouse model in which mutations in the proofreading subunit of the mtDNA polymerase gamma (PolGAD257A) exhibit male infertility due to mitochondrial dysfunction (Kujoth et al., 2005; Trifunovic et al., 2004). The deletion of Mfn1-Mfn2 in the Gc leads to azoospermia in the knock-out mice, suggesting that mitochondrial fusion has an important role in spermatogenesis (Varuzhanyan et al., 2019). Also, the mouse models with a homozygous gene-trap allele of Mff showed reduced sperm count and subfertility, suggesting a role of mitochondrial fission in spermatogenesis (Chen et al., 2015). Recent reports suggest that autophagy plays an important role in post-meiotic spermatids (Shang et al., 2016; Wang et al., 2014). The deletion of Atg7, an important autophagy gene in the primordial germ cells, blocked acrosome biogenesis (Wang et al., 2014).

It is reported that excess mitochondria agglomerate into residual bodies for their phagocytic degradation by Sertoli cells just before the event of spermiation (Dietert, 1966). Some data indirectly indicate prominent mitophagical events in immature and proliferative Sc compared with aging Sc (Eid et al., 2018). Active mitophagy has been observed in neural stem cells before differentiation, aiding in the clearance of faulty mitochondria generated by mitochondrial remodeling (Beckervordersandforth, 2017). We believe that there may be a change in mitophagy-related activity between the proliferation stage of Sc and the differentiated Sc. However, the specific roles of mitophagy-related genes in the development of Sc and on male fertility remain poorly understood. Therefore, we plan to identify and characterize the role of mitophagy-related genes (MRG) in the function of Sc and male fertility to address this gap.

Previously, our group reported that Sc remains in an immature state (more proliferative) in a 5-day-old rat (Bhattacharya et al., 2012). We further reported the differential gene expression profile of rat Sc (Gene Expression Omnibus (GEO) dataset GSE48795) obtained from immature (5 days old) and maturing (12 days old) and matured (60 days old) testes (Gautam et al., 2018). In this study, we used this platform to identify the MRGs that may have an important role in male fertility. From this study, nuclear orphan receptor 1 (*NOR1*), also known as NR4A3, emerged as a key candidate for further investigation due to its regulatory association with Fos. We observed that *NOR1* was up-regulated in the mature Sc, which is associated with robust onset of spermatogenesis. *Nor1*, which belongs to the NR4A subfamily, is a proapoptotic factor (Close et al., 2019). In human beta cells, *Nor1* undergoes mitochondrial translocation upon activation by proinflammatory cytokines and plays a role in mitochondrial fragmentation via mitophagy (Close et al., 2020). In HeLa cells, the loss of *NOR1* causes embryonic lethality and also the partial loss of *Nor1* may also cause embryo lethality at some frequency (DeYoung et al., 2003). The *NOR1* KD in C2C12 myotubes leads to a reduction in the level of PGC-1α, TFAM, and MFN2 (Paez et al., 2023). In the HeLa cells, the *NOR1* upregulation enhanced apoptosis via the mitochondrial-dependent apoptotic pathway (Paez et al., 2023). The Sc-specific knock down of *NOR1* led to compromised spermatogenesis due to enhanced levels of beta-catenin and Smad3 (Shukla et al., 2018). Since the role of *NOR1* has been studied in spermatogenesis in mice models, we generated a transgenic rat model with Sc-specific knockdown of *NOR1* to validate our approach to identifying the MRG gene(s) associated with male fertility. Rat models are physiologically closer to humans as compared to mice models (Pradhan et al., 2016). Our findings revealed that male transgenic rats with *NOR1* knockdown specifically in Sc were infertile, suggesting that the involvement of mitophagy during Sc development and male fertility.

## Results

### Identification of Mitophagy-Related Genes (MRG) during the development of Sc

To investigate the role of mitophagy-related genes in Sc development, we analyzed microarray data from the GSE48795 dataset to identify differentially expressed genes (DEGs). We used an adjusted P value of < 0.05 and a cutoff of |log2(FC)| > 1 to identify significant DEGs. We compared the transcriptomic profile of a 5-day-old Sc with a 12-day-old Sc and identified a total of 1402 DEGs. Similarly, by comparing the RNA-seq data of 5-day-old Sc with 60-day-old Sc, we identified a total of 4612 DEGs. Then, we considered a list of the mitophagy-related differentially expressed genes (MRDEGs) of humans from the available literature to identify MRDEGs in rats. For this, we initially identified the orthologs for these MRDEGs of humans in rats from the NCBI gene. After successful identification of the orthologs, we screened our 99 and 455 MRDEGs for rats in Sc development from 1402 (5 vs 12-day-old Sc) and 4612 (5 vs 60-day-old Sc) DEGs, respectively (**Figure 1A-C and Supplementary Table 1**). Thus, there were a total of 488 mitophagy-related genes in Sc development (MRGSCD) considering both the phases. But, a total of 66 MRGSCDs were identified to be common between the two experimental conditions, i.e. 5d Vs 12 and 5d Vs 60d (**Figure 1D**). Thus, these 66 genes can be considered to be crucial mitophagy-related genes. Functional enrichment analysis of the common 66 MRGSCD showed that GO Biological processes like GO:0022414 – “reproductive process” and GO:0032502-“developmental process” were enriched. Similarly, GO molecular functions like - GO:0005488- “binding” were enriched (**Figure 2**). The reconstruction of a protein interaction network for 66 common MRGSCD would unravel the vital signaling cascade of mitophagy-related genes in Sc development common to both stages. So next, with these 66 MRDEGs, we reconstructed a protein-protein interaction network (PPIN) using STRING v12 (confidence score >0.4) (**Figure 3**). This PPIN consists of 33 nodes and 76 edges. In a PPIN, hub genes are the nodes with the greatest number of edges, or the most connections. Subsequently, using a network analyzer in Cytoscape 3.10.3., we identified 12 hub genes in the PPIN viz. Egfr, Bcl2, Ccl2, Mmp2, Igf1, Fgf7, Apoe, Fos, Cxcl12, Ocln, Dcn, Snca (**Figure 3**). Notably, among these 66 common genes, *NOR1* was highlighted as a key gene of interest due to its significant upregulation in both 12-day and 60-day-old Sc (**Figure 1D and Supplementary Table 1**). Moreover, it was found that *Nor1* was involved in the interactions with only two important hub genes i.e. Bcl2 and Fos in the PPIN. So, we performed further studies on *NOR1* to validate the findings of this analysis.

**Figure 1.**
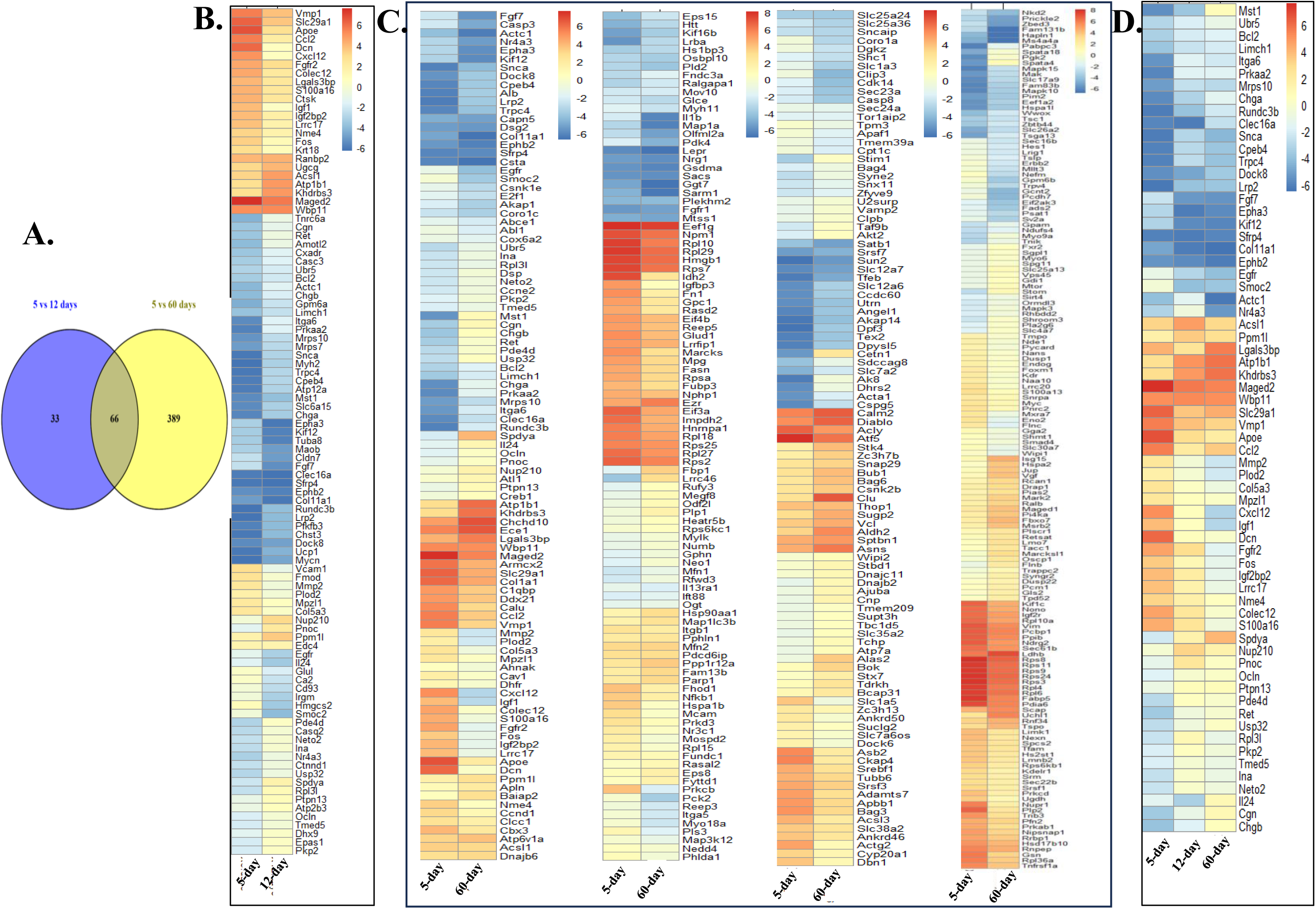
Mitophagy is related to differentially expressed genes for Sc development in rats. **A.** Venn diagram of common MRDEGs among Sc of 12- and 60-day-old rats. Heatmap of average normalized expression of **B.** 99 common MRDEGs in the Sc of 5 and 12-day-old rats. **C.** 455 is common MRDEGs in the Sc of 5 and 60-days old rat. D. 66 is common MRDEGs in the Sc of 5, 12 and 60-day-old rats.

**Figure 2.**
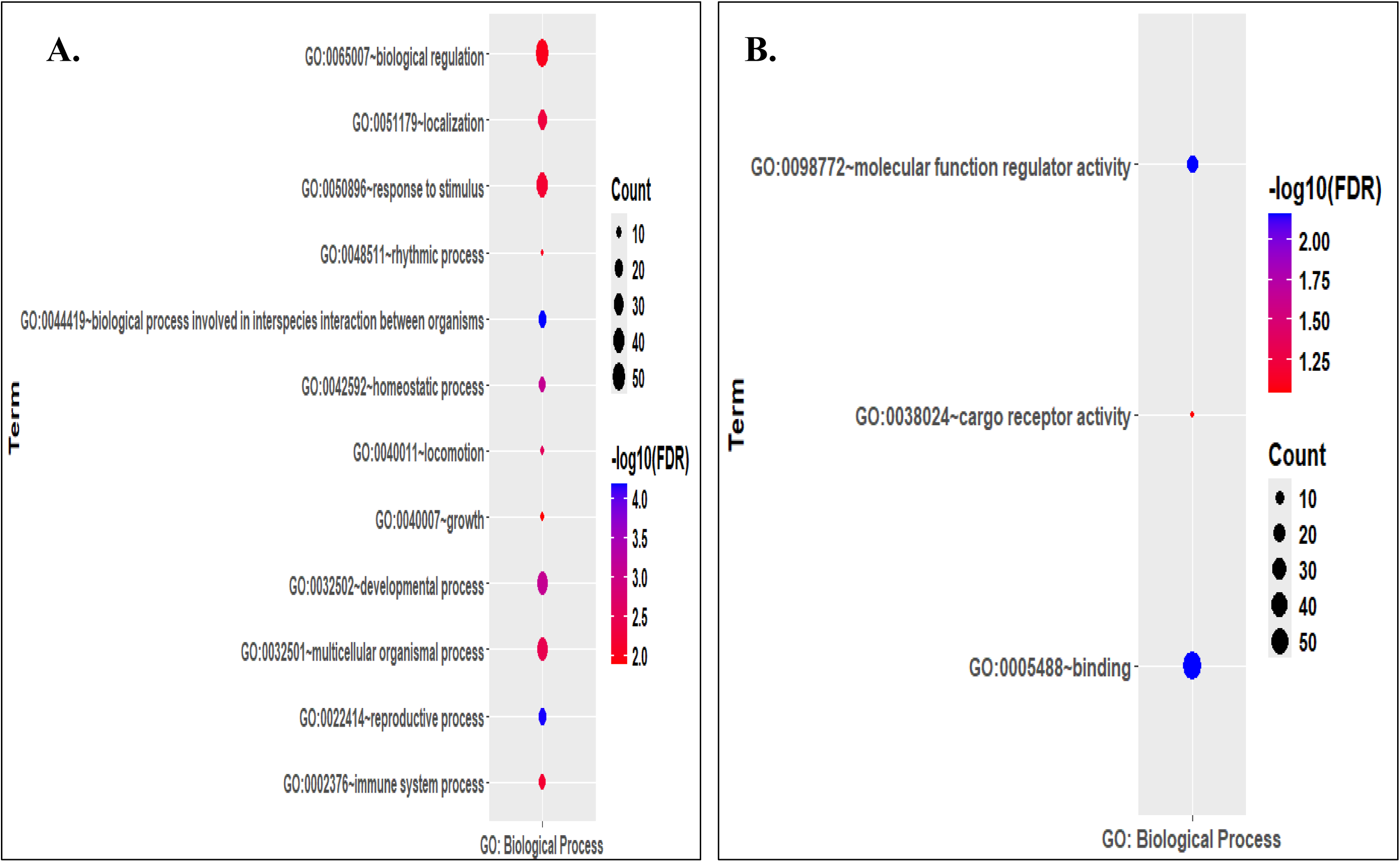
Functional enrichment of 66 common MRDEGs. **A.** GO Biological Process Enrichment **B.** GO Molecular Function enrichment.

**Figure 3.**
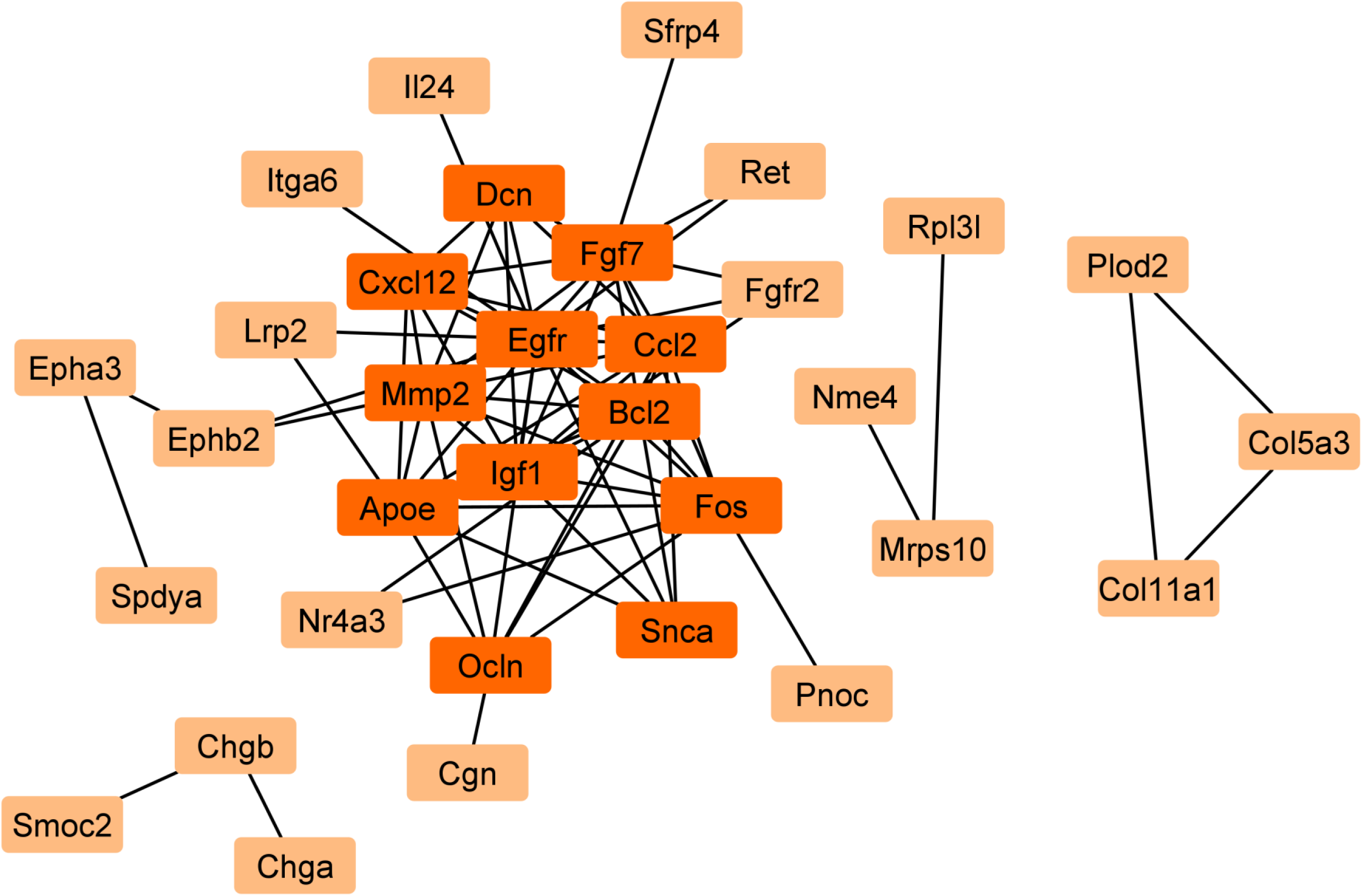
Protein-protein interaction network of mitophagy related to differentially expressed genes. Protein-protein interaction network of 66 mitophagy-related differentially expressed genes (common between Sc of 5 vs 12 and 5 vs 60 old rats) for Sertoli cell development in rats. This network consists of 33 nodes and 76 edges. The orange-colored nodes represent hub genes.

### Validation of the differential expression of *NOR1* in Sc

Since *NOR1* was identified as an important gene related to mitophagy in our analysis, we validated the expression of *NOR1* in the 5-day-old Sc and 12-day-old Sc as it was upregulated in the microarray analysis (**Figure 4A**). These cells were treated with hormones (FSH and T) identical to those of microarray samples. The q-RT-PCR data showed that *NOR1* was significantly (p≤0.05) upregulated in the Sc isolated from the 12-day-old Sc as compared to that of the 5-day-old Sc (n=4) (**Figure 4B**).

**Figure 4.**
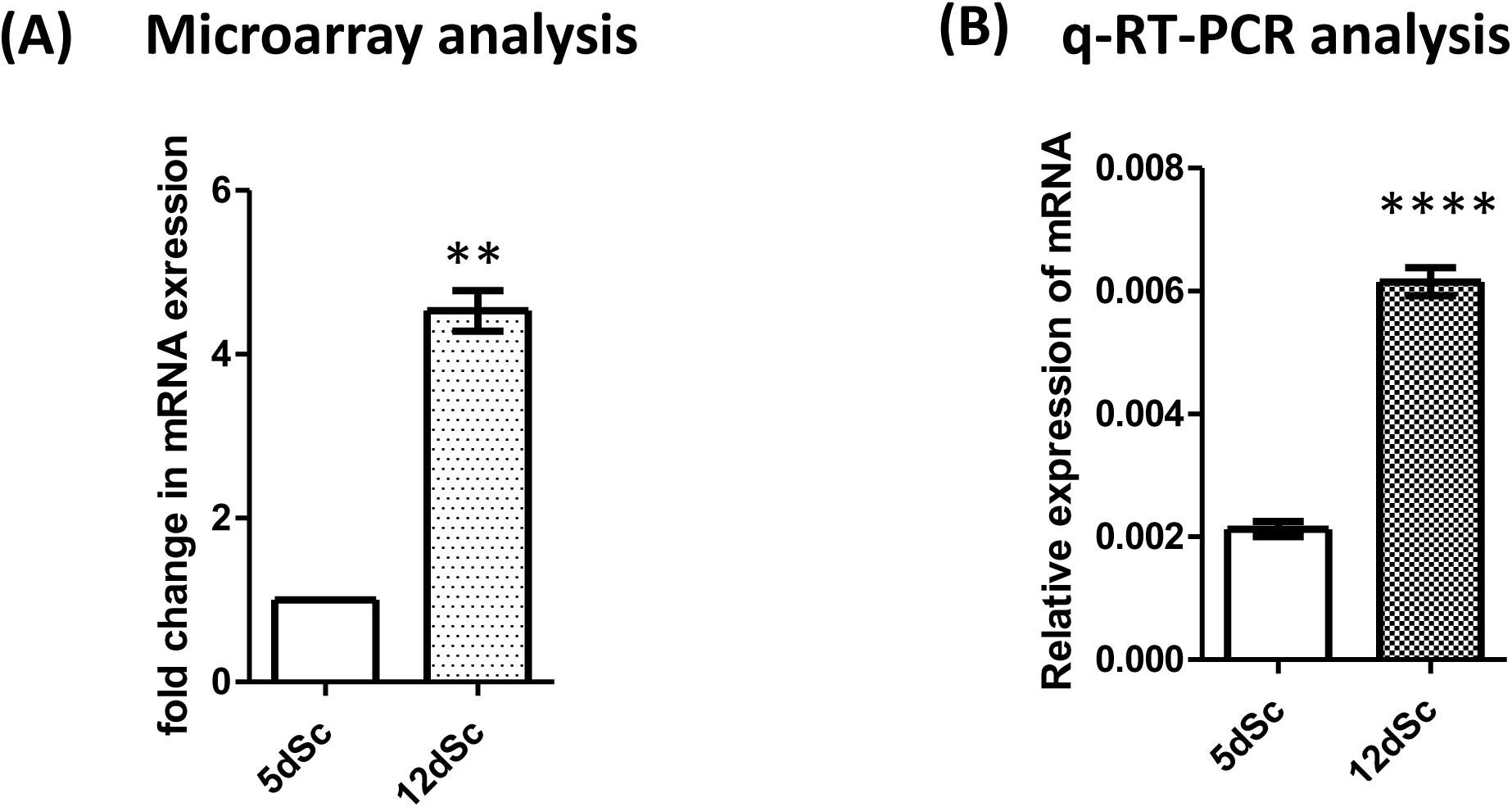
Differential expression of NOR1 in the Sc of the 5d-and 12d old rat. **(A)** NOR1 was upregulated in the Sc of 12d old rat as compared to the Sc of the 12d old rat in the microarray analysis. n=3; paired t-test; **P ≤ 0.01. **(B)** Validation of NOR1 upregulation in the Sc of 12d old rat as compared to Sc of 12d old rat by the q-RT-PCR analysis. Ppia was used as an internal control. N=4, Mann–Whitney test; ****P ≤ 0.0001.

### Generation of transgenic rat with Sc-specific *NOR1* knockdown

Since we identified NOR1 to be upregulated in 12-day-old Sc from the rat microarray data, we generated a transgenic rat with Sc cell-specific knockdown of the NOR1 (Tg rat) using the Pem promoter to determine the functional role of NOR1 in spermatogenesis and on mitophagy. As a control, we generated the transgenic rat with Sc cell-specific knockdown of the LacZ (LacZ control rat). The integration of the transgene was confirmed by slot blot analysis using the GFP probe (**Figure 5 A and B**). Since the Pem promoter drives the expression of shRNA against NOR1 and GFP, we analyzed the testicular section of the Tg rats and age-matched wild type (wt) control rats for the expression of GFP using anti-GFP antibodies to determine the transgene expression. We observed the expression of GFP in the testis of the Tg rat but not in that of the control (**Figure 6**), suggesting that the Pem promoter was active in the Tg rats and able to drive the expression of shRNA. Further, we performed the q-RT-PCR analysis to determine the knock down efficiency of the NOR1 in the testis of Tg rat. We observed that expression levels of NOR1 in the Tg rat were reduced by 47% as compared to the LacZ control rat (**Figure 5C**).

**Figure 5.**
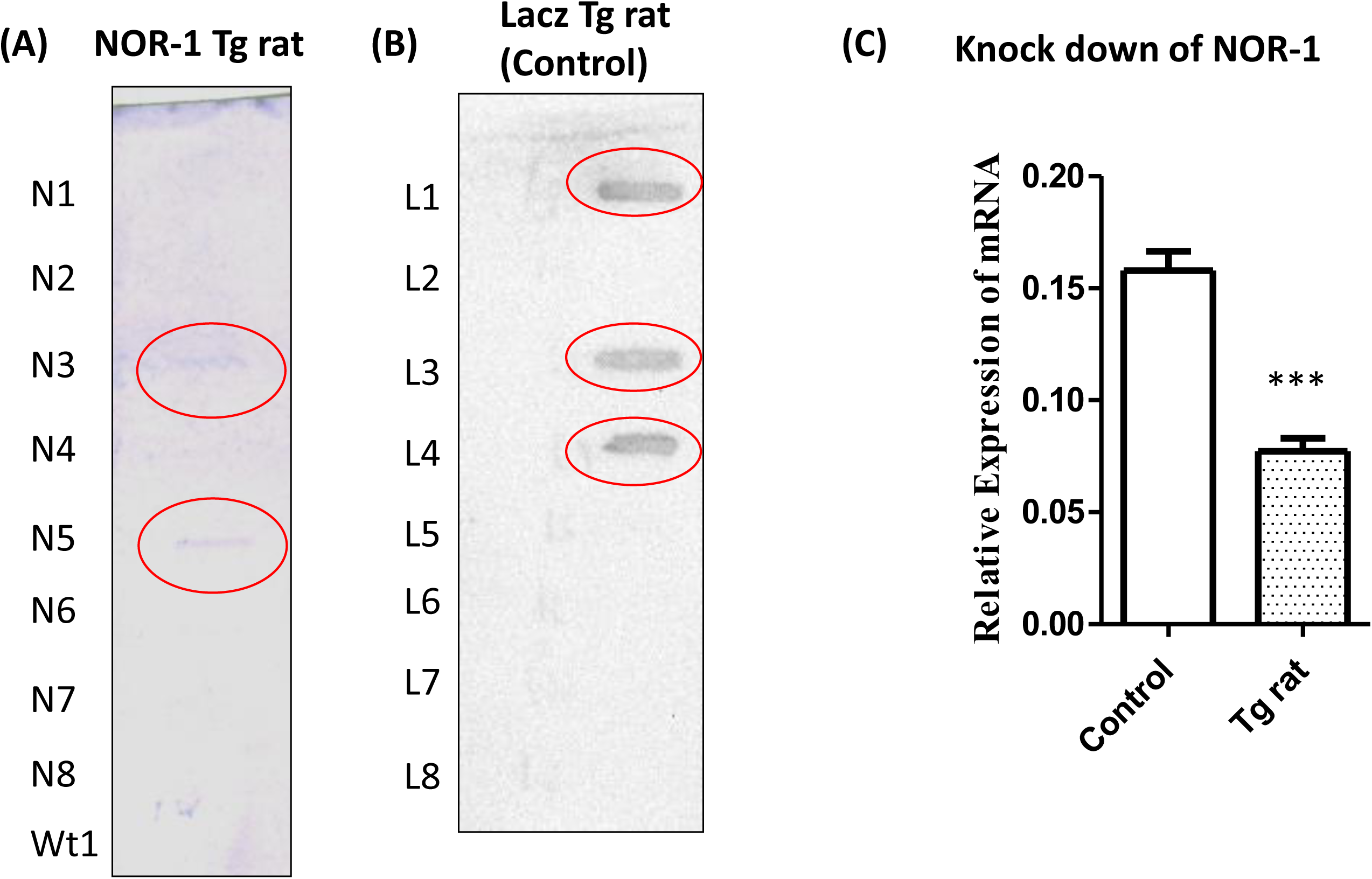
Generation of transgenic rat (Tg rat) with Sc-specific knock-down of NOR1. **(A)** The slot blot analysis of gDNA obtained from the tail of the progenies of the female founder Tg rat (with Sc-specific knock-down of NOR1). Progenies No 3 and No 5 were detected positive for the transgenes. Wt denotes the gDNA obtained from the tail of the wild type rat. N1 to N8 were the pup numbers obtained from the female founder. A part of the GFP was used as a robe to detect the integration of transgene **(B)**. The slot blot analysis of gDNA was obtained from the tail of the progenies of the control Tg rat (with Sc-specific knock down of LacZ). L1 to L8 were the pup numbers obtained from the founder. **(C)** The expression of NOR1 was significantly reduced in the testes of the Tg rats (10 months old) as compared to that of the control as detected by q-RT-PCR. Ppia was used as the internal control. N=3, Mann–Whitney test. ***P ≤ 0.001.

**Figure 6.**
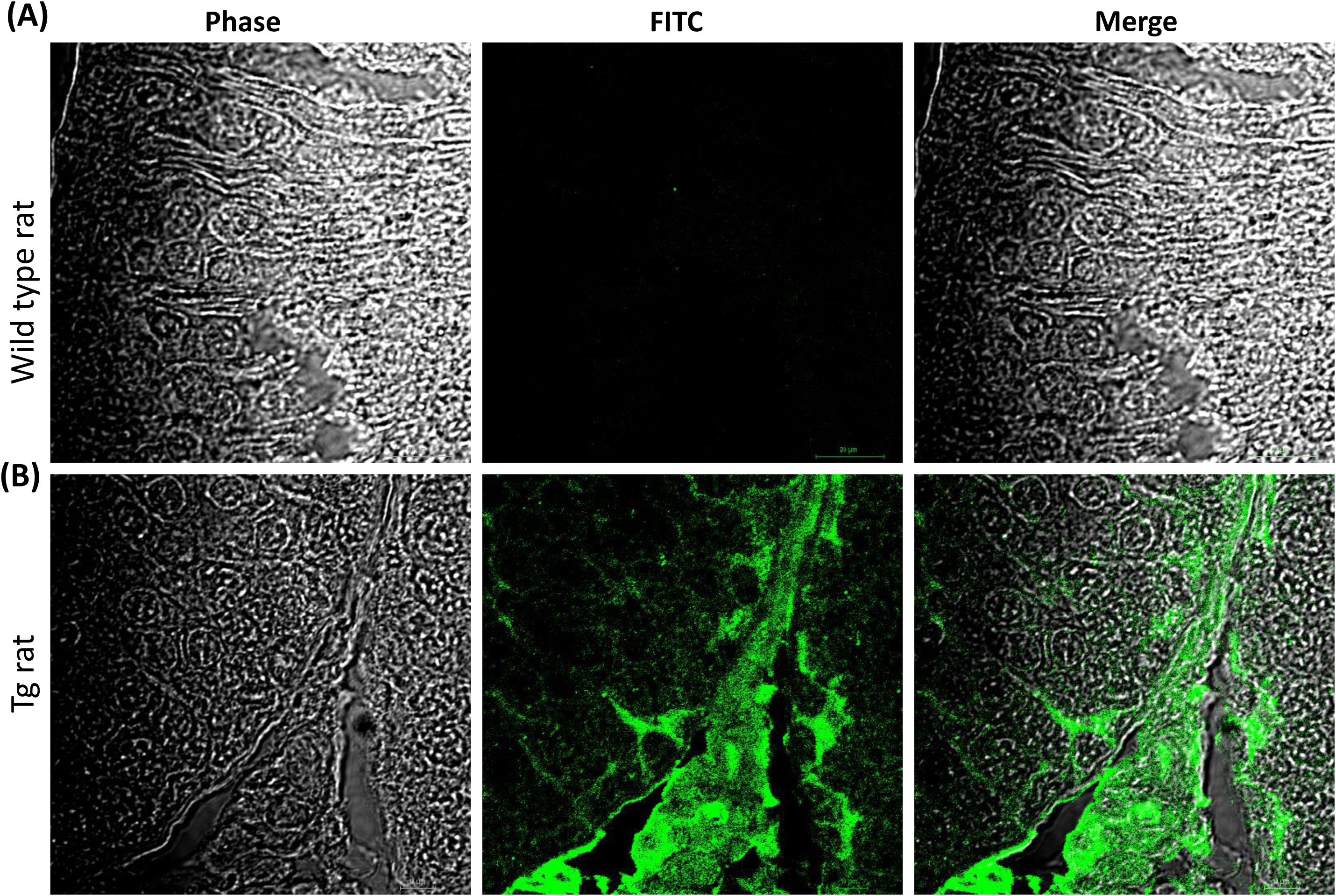
Detection of GFP in the testis of a transgenic rat (Tg rat). **(A)**The expression of GFP was detected in the cross-section of the testis of a Tg rat (10 months old). GFP was used as a marker for the expression of transgene which included shRNA against NOR1 driven by the Pem promoter. **(B)**The cross-section of the testis of the wild type rats was used as the control. N=3.

### Attenuation of spermatogenesis in testes of Tg rats

The Tg rat line could not propagate beyond F1 generation as the positive male pups born out of mating of F1 generation siblings were infertile (**Figure 7A**). On the other hand, the LacZ control rat were maintained beyond the F1 generation. All of our studies were done at F1 generation of shRNA knock down rat as shRNA transgene behaves like dominant-negative alleles of the gene of interest, hence Tg rats can be analyzed in the F1 generation itself without waiting for homozygosity (Gao et al., 2007).

**Figure 7.**
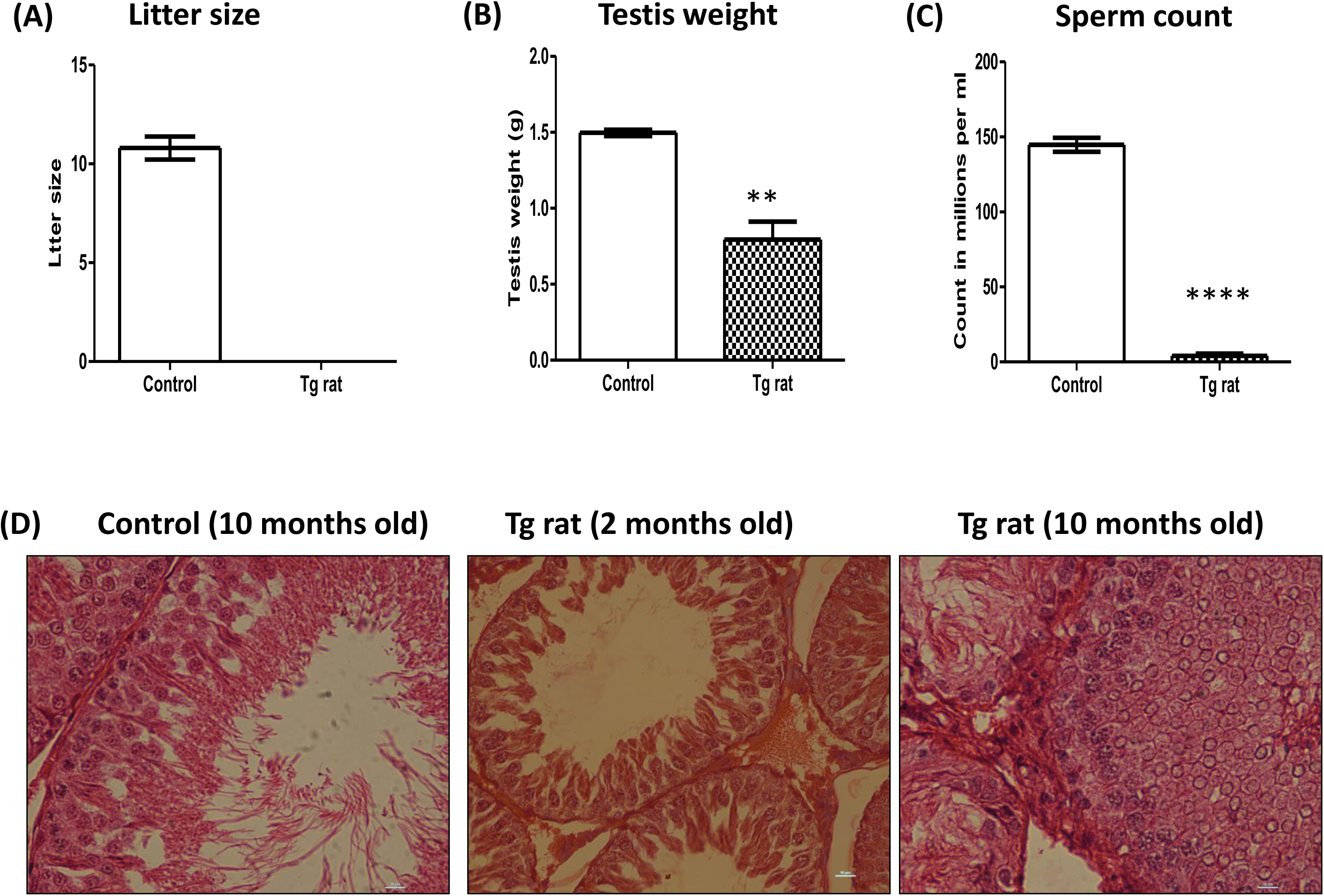
Fertility was inhibited due to impaired spermatogenesis in the Tg rat. **(A)** The litter size of the control and Tg rat. **(B)** The testis weight of the Tg rat was reduced compared to that of the control. N=3, Mann–Whitney test, **P ≤ 0.01. (B) The sperm count of the Tg rat was reduced as compared to that of the control. N=3, Mann–Whitney test, ****P ≤ 0.0001. **(C)** The Hematoxylin and eosin staining of testicular sections obtained from the control rat (10 months old), the Tg rat (2 months old and 10 months old). Spermatogenesis was inhibited in the Tg rat. The images were taken at 40× magnification. N=3.

F1 generation male rat from the Tg rats and LacZ control male rat were analyzed for the defects in reproductive functions. The testis weight of Tg rat (0.79 ± 0.11g) was decreased significantly (p≤0.05) as compared to that of LacZ control rat (1.497±0.02g) (**Figure 7B**). Further, we anlayzed the sperm count from the epididymis of the Tg rats and control. We observed a significant (p≤0.05) reduction in sperm counts in the Tg rat (4 ± 1.52 × 106/ml/epididymis) as compared to their age matched LacZ control rat (144.7 ± 4.667 × 106/ml /epididymis) (**Figure 7C**). Although total sperm count was found to be drastically reduced, more than 80% of the sperm were found to be motile in Tg rats.

The qualitative histopathological evaluation of the testis of the Tg rats (2 months old and 10 months old) and LacZ control rat (10 months old) were performed to determine the effect of NOR1 knock down. We observed that LacZ control rat showed a normal tubular structure whereas the Tg rats showed the sloughing of spermatogenic cells in majority of the tubules (**Figure 7D**). Moreover, multiple vacuoles were observed in this seminiferous epithelium of the Tg rat suggesting that spermatogenesis was impaired due to the decline of NOR1.

## Discussion

Our study demonstrated that there is a change in mitophagy-related genes during the development of Sc. We identified 12 mitophagy-related hub genes - Egfr, Bcl2, Ccl2, Mmp2, Igf1, Fgf7, Apoe, Fos, Cxcl12, Ocln, Dcn, and Snca, which are differentially expressed in our selected age group i.e. 5 days old Sc, 12 days old Sc and 60 days old Sc. To determine the importance of our analysis of finding the mitophagy-related genes, we performed a functional genomics study. We identified NOR1 as a key gene related to mitophagy which is involved in the interactions with two important hub genes i.e. Bcl2 and Fos. The expression of NOR1 was upregulated in the 12-day-old Sc as compared to the 5-day-old Sc, and we validated the differential expression of NOR1. To determine its effect on mitophagy and on spermatogenesis, we generated a Tg rat with reduced expression of NOR1 specifically in the Sc of the pubertal rats. The male Tg rats were infertile due to sperm count and showed abnormalities in the testicular architecture. Overall, our data suggest that NOR1 is an important mitophagy-related gene and plays a crucial role in spermatogenesis.

From the differential gene expression profile list, our main focus was to identify the hub genes which may regulate the mitophagy of Sc during development. Epidermal growth factor receptor (EGFR) is a receptor tyrosine kinase (Sharma et al., 2003). The mitochondrial translocation of EGFR may enhance cancer metastasis by modulating the mitochondrial dynamics by inhibiting the polymerization of Mfn1 (Che et al., 2015). The male transgenic mice overexpressing human EGF are infertile (Wong et al., 2000) whereas knock out mice lacking the growth factor of EGFR - amphiregulin (AREG), betacellulin (BTC), epidermal growth factor (EGF), epigen (EPGN), epiregulin (EREG), heparin-binding EGF-like growth factor (HBEGF), and transforming growth factor alpha (TGFA) are fertile (Schneider et al., 2010). In our study, we observed a down-regulation of EGFR during the differentiation of Sc i.e. EGFR was upregulated at 5 days old Sc and down regulated to 12 days old Sc and its level declined further to 60 days old Sc.

The BCL2 apoptosis regulator is a 26-kDa protein involved in regulating cell death and located at the contact sites between the inner and outer mitochondrial membranes (Hockenbery et al., 1990; Strasser et al., 1990). Bcl-2-deficient mice have normal spermatogenesis (Russell et al., 2002). Bcl-2 suppressed mitophagy through inhibition of Parkin translocation to depolarized mitochondria (Hollville et al., 2014). We observed the upregulation of Bcl2 in the 12-day-old Sc and 60-day-old Sc as compared to the 5-day-old Sc. The C-C Chemokine Ligand 2 (CCL2) is a chemokine and plays an important role in the process of autophagy and mitochondrial biogenesis (Murphy and Hartley, 2018; O’Connor et al., 2015). The Ccl2 KO mice are viable and fertile (Kurihara et al., 1997; Lu et al., 1998). The overexpression of CCl2 in mice leads to an increase in mitochondria due to less degradation of mitochondria in the liver (Luciano-Mateo et al., 2020). We observed a reduction in the level of Ccl2 in the 12-day-old Sc and 60-day-old Sc as compared to the 5-day-old Sc.

Matrix metalloproteinase-2 (MMP-2) is localized to the mitochondria-associated membrane of the heart (Hughes et al., 2013). It proteolyzes mitofusin-2 during myocardial ischemia (Bassiouni et al., 2023). The MMP-2 knock out mice are fertile (Itoh et al., 1997). Previous studies suggested that FSH upregulated the expression of MMP-2 in Sc (Longin and Le Magueresse-Battistoni, 2002). It plays a role in basement membrane remodeling during the release of differentiated germ cells from the basement membrane (Saengsoi et al., 2011). We observed a decline in the level of Mmp2 in the 12-day-old Sc and 60-day-old Sc as compared to 5-day-old Sc. Insulin-like Growth Factor I (Igf1) prevents aging by the activation of mitophagy via activation of Nrf2/Sirt3 signaling (Hou et al., 2020). Previous studies reported that IGF-1 signaling regulates the dynamics of mitochondria via the GSK-3β-Nrf2-BNIP3 pathway (Riis et al., 2020). The IGF-1-KO mice were fertile (Yakar et al., 2001). We observed a decline in the level of Igf1 in the 12-day-old Sc and 60-day-old Sc as compared to 5-day-old Sc.

Fibroblast growth factor 7 (FGF7) is the seventh member of the FGF family and protects osteoblasts against oxidative damage through targeting mitochondria and upregulate the expression of MFN1, MFN2, and LONP1, which are related to mitochondrial fusion and elongation (Liu et al., 2024). It also improves oxidative stress via PI3Kα/AKT-mediated regulation of Nrf2 and HXK2 and alleviates myocardial infarction (Mei et al., 2022). The Fgf7 knockout mice are fertile (Guo et al. 1996). We observed a decline in the level of Fgf7 in the 12-day-old Sc and 60-day-old Sc as compared to 5-day-old Sc.

ApoE is upregulated in various models of mitochondrial respiratory chain dysfunction (Wynne et al., 2023). The mice homozygous for the apoE mutation are fertile. We observed a decline in the level of Apoe in the 12-day-old Sc and 60-day-old Sc as compared to the 5-day-old Sc. The mice deficient in FOS are fertile, although they have reduced placental and fetal weight (Hu et al., 1994). cFos signaling may promote mitophagy. We observed a decline in the level of Fos in the 12-day-old Sc and 60-day-old Sc as compared to 5-day-old Sc. Stromal cell-derived factor 1α (SDF-1), also known as CXCL12, is an important member of the CXC family of chemokines. The CXCL12 knocks out mice that are fertile (Greenbaum et al., 2013). The CXCL12/CXCR4 axis supports mitochondrial trafficking in the tumor myeloma microenvironment (Giallongo et al., 2022). Previous reports suggest the role of the CXCR4/CXCL12 axis in the prevention of ROS elevation, apoptosis and DNA damage in HSCs (Zhang et al., 2016). We observed a decline in the level of Cxcl12 in the 12-day-old Sc and 60-day-old Sc as compared to the 5-day-old Sc.

Occludin (Ocln) is an epithelial tight junction protein and the mislocalization of ocln leads to apoptosis through the extrinsic pathway (Beeman et al., 2009). The Ocln knockout male mice are infertile (Saitou et al., 2000). We observed an increase in the level of Ocln in the 12-day-old Sc and 60-day-old Sc as compared to the 5-day-old Sc. Decorin (Dcn) promotes mitophagy via mitostatin by interacting with the MET receptor tyrosine kinase (Neill et al., 2014). The decorin that knocks out mice is fertile (Danielson et al., 1997). We observed a decline in the level of Dcn in the 12-day-old Sc and 60-day-old Sc as compared to the 5-day-old Sc. The high levels of α-Synuclein (Snca) disrupt mitochondria and impair respiration (Pathak et al., 2011). The supra-physiologic levels of Snca promote excessive mitophagy (Choubey et al., 2011). On the other hand, the reduced level of Snca protects against mitochondrial toxins (Pathak et al., 2011). We observed an up-regulation in the level of Scna in the 12-day-old Sc and 60-day-old Sc as compared to 5-day-old Sc.

The identification of these differentially expressed hub genes related to mitophagy suggests that the developmental changes in the Sc are associated with mitophagy. To confirm this further, we selected NOR1 for further study as it is upregulated in the 12-day-old Sc and 60-day-old Sc as compared to 5-day-old Sc. Also, NOR1 is associated with two hub genes - Bcl2 and Fos. Bcl2 was also upregulated in the 12-day-old Sc and 60-day-old Sc as compared to 5-day-old Sc and Bcl-2 suppressed mitophagy (Hollville et al., 2014). Nor1 knockout mice generated by deleting a portion (37%) of its N-terminal transactivation domain and the first zinc finger domain die during embryonic stages (DeYoung et al., 2003). In contrast to this, mutation of the Nor1 gene locus leads to viable homozygous animals with only a minor problem in inner ear development (Ponnio et al., 2002). Recently, transgenic mice with Sc-specific decline in NOR1 led to germ cell apoptosis and reduced fertility (Shukla et al., 2018), suggesting that NOR1 may have a role in spermatogenesis. Therefore, we generated the Tg rat with Sc-specific reduction of NOR1 to determine the validity of our analysis of determining mitophagy-related genes in Sc. We observed that the Tg rats with Sc-specific decline in NOR1 were infertile due to reduced sperm count and abnormal testicular structure. These findings are in agreement with previous reports (Shukla et al., 2018) on NOR1, suggesting that functional analysis of NOR1 in vivo and identification of mitophagy-related hub genes are reliable.

The limitation of this study is that we did not perform any studies on the functional aspect of mitophagy in these Tg rats due to the lower number of Tg rats generated by these lines. Further study will be needed to determine the effect of NOR1 on mitochondrial structure and function in these Tg rats.

In conclusion, we identified 12 key hub genes related to mitophagy which were differentially expressed during the development of Sc. Moreover, we generated a transgenic rat with Sc-specific decline in NOR1which was infertile due to defects in spermatogenesis.

## Materials and Methods

### Animals and reagents

Wistar rats (Rattus norvegicus) were obtained from the Small Animal Facility of the National Institute of Immunology (New Delhi, India). All animals were housed and used as per the national guidelines provided by the Committee for the Purpose of Control and Supervision of Experiments on Animals. Protocols for the experiments were approved by the Institutional Animal Ethics Committee (IAEC) of National Institute of Immunology (New Delhi, India) and the Committee for the Purpose of Control and Supervision of Experiments on Animals (CPCSEA). All other reagents, unless stated otherwise, were procured from Sigma Chemical (St. Louis, MO).

### Screening of hub genes for Sertoli cell development in Rat

We identified differentially expressed genes (DEGs) by analyzing microarray data from the GSE48795 dataset (Gautam et al., 2014) in order to examine the function of mitophagy-related genes in Sertoli cell development. The Bioconductor project’s basic packages in the NCBI Gene Expression Omnibus (GEO) database, which were based on limmav3.26.8, were utilized for data retrieval, logarithmic transformation, and quantile normalization of the data (Ritchie et al., 2015). A cut-off of adjusted P value < 0.05 and |log2(FC)| > 1 was considered to identify significant DEGs. The orthologs for humans in rats were identified from NCBI’s genes (Sayers et al., 2022). Then, to reconstruct the PPIN for Sertoli cell development, we considered both physical and functional interactions from STRING v12 (Szklarczyk et al., 2023). The default cut off i.e. confidence score >0.4 was considered for the reconstruction of the PPIN. To identify hub genes, we used a network analyzer in Cytoscape 3.10.3 (Shannon et al., 2003) to calculate network degree centrality. A network degree centrality >average of total degree count of the nodes i.e > 4.5 was considered as a cut-off for identifying hub genes.

### Functional enrichment analysis

We used DAVID v2023q4 (Dennis et al., 2003) for GO Biological Process and GO Molecular Function enrichment of the genes.

### Differential expression analysis of *NOR1* in 5 days and 12 days old rat Sc culture

We identified NOR1 as a candidate gene from the analysis. The differential expression of NOR1 from this analysis was validated by q-RT-PCR analysis in the three separate sets of Sc cultures of 5-day and 12-day-old rat testis. The Sc were isolated and cultured, treated with FSH and Testosterone in combination on day 4 of culture as reported by us previously (Bhattacharya et al., 2011; Bhattacharya et al., 2017; Gautam et al., 2014). Further, the total RNAs were extracted from these groups of cells as described previously (Pradhan et al., 2019; Pradhan et al., 2020).

### Plasmids and Cloning

The knock down of NOR1 using shRNA in the Sc of the prepubertal rats were performed as described previously (Basu et al., 2018; Mandal et al., 2018; Shukla et al., 2018). shRNA was designed by using online tools available at Dharmacon (http://www.thermoscientificbio.com/design-center/) and Clontech (http://www.clontech.com/US/Support/xxclt_onlineToolsLoad.jsp?citemId= http://bioinfo.clontech.com/rnaidesigner/oligoDesigner.do?overhangs=on&restrictionSite=on§ion=16260&xxheight=750). shRNA sequences were verified for target specificity by BLAT (http://genome.ucsc.edu/cgi-bin/hgBlat?command=start) and BLAST analysis (http://www.ebi.ac.uk/Tools/sss/ncbiblast/). The shRNA against NOR1 and bacterial LacZ (as a control) were designed and cloned under the Pem promoter. Pem promoter is reported to drive shRNA expression specifically in pubertal Sertoli cells (Lindsey and Wilkinson, 1996; Rao and Wilkinson, 2006). The sequence for shRNA for NOR1 was *TCGAGAGACAAGAGACGTCGAAATTTCAAGAGAATTTCGACGTCTCTTGTCTTTTTTTACCGGTCCGC* and the sequence for shRNA for LacZ was *TCGAGGCATCGAGCTGGATAATAATTCAAGAGATTATTATCCAGCTCGATGCTTTTTTACCGGTCCGC*.

### Generation of transgenic (Tg) rats with reduced *NOR1* in Sc

The construct was linearized with Stu I, and was used to generate a transgenic rat (Tg) as reported by us previously (Pradhan et al., 2016; Pradhan et al., 2019; Pradhan et al., 2020). Briefly, the linearized construct (a total of 30 µg DNA with a concentration of 1 µg/µl) was injected into the testis of a 38-day-old Wistar rat, and electroporated using 8 square 90 V electric pulses in alternating directions with a time constant of 0.05 second and an inter-pulse interval of ∼1 second via an electric pulse generator. The electroporated male rats were mated with wild-type female rats 60 days post electroporation. Born pups were screened by PCR using genomic DNA obtained from their tail biopsies and those positive for the transgene were regarded as F1 generation of transgenic (Tg) rats.

The integration of transgene in the transgenic rats was confirmed by slot blot analysis as described previously (Basu et al., 2018; Mandal et al., 2018; Shukla et al., 2018). For slot blot analysis, around 2µg of gDNA isolated from the tail biopsies were denatured at 950C for 10 min and blotted on to Hybond N+ (Amersham Pharmacia Biotech, England) in a slot blot apparatus (Cleaver Scientific Co., England) under vacuum. The membrane was UV cross-linked (Church and Gilbert, 1984) at 12 x 104 µJ/cm2 energy in a CL-1000 Ultraviolet crosslinker (UVP, Upland, CA, USA). The non-radioactive DIG probe was used to detect the transgene. The probe was prepared by amplifying a fragment that contained part of the GFP (630 bp) using the primers: GFPF: GACGTAAACGGCCACAAGTT, GFPR: GGCGGTCACGAACTCCAG and during PCR, the probe was labeled with DIG. The membranes were pre-hybridized for 2h at 37°C in hybridization solution without labeled probe and then hybridized separately at 37°C with probe specific for GFP for 16 h. The membranes were washed twice, for 5 min each, at room temperature in 2Х saline sodium citrate buffer and 0.1% SDS. This was followed by another washing for 15 min at 60°C (in 0.1Х saline sodium citrate buffer and 0.1% SDS). Detection was performed by using the DIG labeling and BCIP/NBT substrate according to the manufacturer’s recommendation (DIG system user’s guide for filter hybridization, Roche, Germany).

### Validation of knock down of *NOR1* by q-RT-PCR

Total RNA was extracted from the whole testicular extract of control and Tg rats of 10 months old (n=3). q-RT-PCR amplifications were performed to detect expression of *NOR1* and *ppia* (loading control) using the RealplexS (Eppendorf) as described previously (Pradhan et al., 2020). Primers for *NOR1* were *NF: GCTTGTCTCTGCACCATTCA; NR: TGAGCTAGGCCTCGAAGGTA.* Primers for *ppia* were *PF: ATGGTCAACCCCACCGTGT; PR: TCTGCTGTCTTTGGAACTTTGTCT.* The expressions of mRNA of the target genes were evaluated by the efficiency corrected 2−ΔΔCT method.

### Immunofluorescence microscopy

The testis of Tg (10 months old) and age-matched WT rat were collected, and fixed in Bouins solution, and IF was performed as described previously (Pradhan et al., 2016). To stain for transgene GFP, a primary Rabbit polyclonal anti-GFP antibody was used at a dilution of 1:250 and incubated for 2 h at room temperature. A Goat anti-Rabbit IgG conjugated with Alexa 488 (Thermoscientific, USA) at a dilution of 1:1000 was used as the secondary antibody.

### Tissue histology

Tissue histology was performed as described previously (Pradhan et al., 2020). Briefly, the testicular tissue samples of rats were fixed in Bouins solution at room temperature for 24 hours. Dehydration of tissues was done in a series of ascending concentrations of ethanol for 1 hour in each grade of ethanol. The tissues were embedded in paraffin, and 4 µm sections were cut. Sections were stained with hematoxylin and eosin, and were examined to evaluate the status of spermatogenesis in the control (10 months old) and Tg rat (2 months and 10 months old).

### Fertility analyses of *NOR1* knock down rat

Testes weight of NOR1 knock down (10 months old) and LacZ shRNA used as control males were recorded. The numbers of epididymal spermatozoa were analyzed as described previously (Pradhan et al., 2020). Total numbers of sperm present in each epididymis were counted after releasing the sperms in 1ml of 1× PBS by mincing the epididymis. Motility of at least 200 epididymal spermatozoa per field was assessed under light microscope.

### Statistical analysis

For the validation of differential expression of *NOR1* by q-RT-PCR analysis were analyzed using Mann–Whitney tests. For fertility studies, at least three or more sets of observations were analyzed by Mann–Whitney tests. A value of *P* ≤ 0.05 was considered as significant. All statistical analyses were performed using GraphPad Prism v. 5.01 (GraphPad Software, La Jolla, CA, USA).

## Data availability

All the data were provided as the supplementary dataset in the manuscript.

## Conflicts of Interest

The authors declare that they have no competing interests.

## Funding

We are grateful to Department of Biotechnology, Govt. of India, for providing the financial assistance under grants BT/PR11313/AAQ/01/376/2008, BT/HRD/35/01/01/2010 (TATA Innovation Award) and BT/PR10805/AAQ/1/576/2013. BSP was supported the Polish National Science Centre grant (2020/39/D/NZ5/02004).

## Author Contributions

Experiments were conceived and designed by B.S.P., and S.S.M. Experiments were performed by B.S.P., D.D. Data of the manuscript was analyzed by B.S.P, H.S., I.B. Primary draft of the manuscript was written BSP; and reviewed by B.S.P. D.D., H.S and IB. All authors have read and agreed to the published version of the manuscript.

## Acknowledgments

We are thankful to all the staff of the Small Animal Facility. Thanks are due to Ram Singh, Dharamvir Singh and Birendar Roy for the technical assistance. We are grateful to the Director, NII for valuable support.

---

**Supplementary File 1. List of MRDEGs for Sc of 12- and 60-day-old rats**.

## Notes

### Competing Interest Statement

The authors have declared no competing interest.

